# The microRNA miR-132 is a key regulator of lymphatic vascular remodelling

**DOI:** 10.1101/2021.12.22.473780

**Authors:** Valeria Arcucci, Musarat Ishaq, Sally Roufail, B. Kate Dredge, Andrew G. Bert, Emily Hackett-Jones, Ruofei Liu, Katherine A. Pillman, Stephen B. Fox, Steven A. Stacker, Gregory J. Goodall, Marc G. Achen

**Affiliations:** Tumour Angiogenesis and Microenvironment Program, Peter MacCallum Cancer Centre, 305 Grattan St., Melbourne, Victoria 3000, Australia; Sir Peter MacCallum Department of Oncology, University of Melbourne, Parkville, Victoria 3010, Australia; O’Brien Institute Department, St Vincent’s Institute of Medical Research, 9 Princes Street, Fitzroy, Victoria 3065, Australia; Department of Medicine, St Vincent’s Hospital, University of Melbourne, Fitzroy, Victoria 3065, Australia; Centre for Cancer Biology, SA Pathology and University of South Australia, Adelaide, SA, Australia; Department of Pathology, Peter MacCallum Cancer Centre, 305 Grattan St., Melbourne, Victoria 3000, Australia; Department of Surgery, Royal Melbourne Hospital, The University of Melbourne, Parkville, Victoria 3050, Australia; Discipline of Medicine, University of Adelaide, Adelaide, SA, Australia; School of Molecular and Biomedical Science, University of Adelaide, Adelaide, SA, Australia

## Abstract

Lymphangiogenesis (growth of new lymphatic vessels), and lymphatic remodelling more broadly, are important for disease progression in cancer, lymphedema and the pulmonary disease lymphangioleiomyomatosis. Multiple molecular pathways which signal for aspects of lymphangiogenesis are known but little is understood about their co-ordinate regulation in lymphatic endothelial cells (LECs). Small RNA molecules co-ordinately regulate complex biological processes, but knowledge about their involvement in lymphangiogenesis is limited. Here we used high-throughput small RNA sequencing of LECs to identify microRNAs (miRs) regulating lymphatic remodelling driven by the lymphangiogenic growth factors VEGF-C and VEGF-D. We identified miR-132 as up-regulated by both growth factors, and demonstrated that inhibiting miR-132 in LECs *in vitro* blocked cell proliferation and tube formation, key steps in lymphangiogenesis. We showed that miR-132 is expressed in human LECs *in vivo* in the lymphatics of human breast tumours expressing VEGF-D. Importantly, we demonstrated that inhibiting miR-132 *in vivo* blocked many aspects of lymphangiogenesis in mice. Finally, we identified mRNAs regulated by miR-132 in LECs, by sequencing after RNA-protein cross-linking and Argonaute immunoprecipitation, which demonstrated how miR-132 co-ordinately regulates signalling pathways in lymphangiogenesis. This study shows miR-132 is a critical regulator of lymphangiogenesis and a potential target for therapeutically manipulating lymphatic remodelling in disease.

## INTRODUCTION

The lymphatic vasculature is a critical feature of many tissues and organs which is required for maintenance of tissue fluid homeostasis, immune function and absorption of fats and fat-soluble vitamins (1). It consists of multiple types of lymphatic vessels: initial lymphatics, which take up tissue fluid; lymphatic pre-collectors and collectors which drain lymph fluid from tissues; lymphatic ducts which return lymph to the venous circulation (2). Lymphatic vessels can undergo remodelling involving modifications in their shape, size and molecular features. A prominent form of lymphatic remodelling is lymphangiogenesis which is the formation of new lymphatic vessels from pre-existing lymphatics. Lymphangiogenesis involves proliferation, sprouting, migration and tube formation by lymphatic endothelial cells (LECs) (3). Remodelling of the lymphatic vasculature occurs during embryonic development and in several pathological conditions such as cancer, inflammation, infection, lymphedema, lymphangiectasia and lymphangioleiomyomatosis (LAM) (4–7). Lymphatic remodelling can be important for disease progression, e.g. lymphangiogenesis in a primary tumour can facilitate the metastatic spread of cancer (3,8,9).

From a molecular perspective, the remodelling of lymphatic vessels can be driven by soluble growth factors which bind cognate receptors on LECs and activate molecular pathways that drive cell proliferation, migration and tube formation. VEGF-C and VEGF-D are two such growth factors which induce lymphangiogenesis by activating VEGFR-3, a receptor tyrosine kinase localized on the surface of LECs (10–13). VEGF-C and VEGF-D are the most extensively studied lymphangiogenic factors that have been shown to induce lymphatic remodelling in embryonic development, cancer and other disease settings (6, 14). The activation of VEGFR-3 on LECs by VEGF-C or VEGF-D leads to activation of a variety of downstream signalling pathways including the PI3K-AKT pathway (15), important for cell proliferation, and pathways which play a role in tube formation by LECs (16). Furthermore, other signalling pathways important in lymphangiogenesis have recently been identified by a genome-wide functional screen based on primary human LECs (17). However, the means by which such signalling pathways are co-ordinately regulated in LECs to drive growth and remodelling of lymphatics is not well understood.

MicroRNAs (also known as miRNAs or miRs) are small RNA molecules which mediate post-transcriptional regulation by targeting mRNAs as part of the Argonaute-containing RISC complex; specificity is achieved based on complementarity between the “seed” region of the miRNA, and a target region which is usually in the 3’-untranslated region (UTR) of the regulated mRNA (18). This interaction leads to decreased gene expression due to mRNA cleavage, mRNA destabilization, translational repression or combinations of these effects (19). So far, thousands of miRNAs have been identified among many animal species, and they have been found to regulate a vast array of complex biological processes by controlling molecular pathways. The increased availability of data, generated from high throughput technologies analysing miRNA targets, has revealed that a single miRNA can control a molecular pathway, or networks of pathways, in a cell by co-ordinated global targeting of multiple genes in one or multiple pathways. Moreover, multiple miRNAs can interact in reciprocal feedback to control pathways and networks of pathways (20). Notably, experimental evidence has demonstrated that miRNA expression and function can be dysregulated in pathological settings including cancer, inflammation, infection and LAM (21–24). Although some miRNAs have been proposed to be involved in lymphangiogenesis (25–27), there have been no comprehensive analyses of the role played by these important regulatory molecules in lymphatic remodelling.

Here, we have comprehensively identified miRNAs whose expression in LECs is regulated by VEGF-C and/or VEGF-D, with a view to discovering miRNAs which are mechanistically important for lymphangiogenesis. We showed that these growth factors modulate the levels of over 60 miRNAs in LECs, including miR-132 which is significantly up-regulated by both VEGF-C and VEGF-D. We demonstrated that miR-132 is critical for proliferation and tube formation of LECs, and for lymphangiogenesis *in vivo*. We then identified the miR-132-mRNA interactome in LECs which highlighted how miR-132 co-ordinately regulates multiple signalling pathways involved in the processes of LEC proliferation, tube formation and sprouting which are required for lymphatic remodelling.

## RESULTS

### miRNA levels in LECs are profoundly altered in response to lymphangiogenic growth factors

We are interested in understanding how molecular signalling pathways are co-ordinately regulated to drive lymphangiogenesis and lymphatic remodelling. Given that miRNAs can modulate the activity of multiple signalling pathways in a co-ordinated way, and thereby globally regulate molecular networks and complex biological processes, we analysed LECs for changes in miRNA levels in response to VEGF-C and VEGF-D. Human primary LECs were treated *in vitro* with mature human forms of VEGF-C or VEGF-D for 3 h, a time-point previously shown to be sufficient for up- or down-regulation of miRNAs to occur in response to extra-cellular signalling molecules such as growth factors (28). Changes in the levels of all miRNAs were analysed by small RNA Sequencing (small RNA-Seq) (Figure 1A). This revealed either up- or down-regulation of 63 miRNAs in response to these growth factors compared with a negative, vehicle-only, control (>10 reads per million, >1.5-fold change in miRNA level). Among those 63 miRNAs, 12 responded to treatment with either VEGF-C or VEGF-D, 11 to VEGF-C only and 40 to VEGF-D only (Figure 1A-C). The miRNA that was changed most profoundly in abundance in response to VEGF-C was miR-132-3p (referred to here as miR-132) which was increased 7.3-fold, and this miR was also up-regulated 1.6-fold by VEGF-D, compared with LECs which were not treated with these growth factors. Notably, miR-132 has been studied in several biological contexts including tumour growth and angiogenesis (28, 29). Given its significant degree of regulation by both VEGF-C and VEGF-D, and its previously identified role in blood vessel biology, miR-132 was chosen for validation and further analysis, in order to determine if it plays a functionally important role in lymphangiogenesis and lymphatic remodelling.

**Figure 1.**
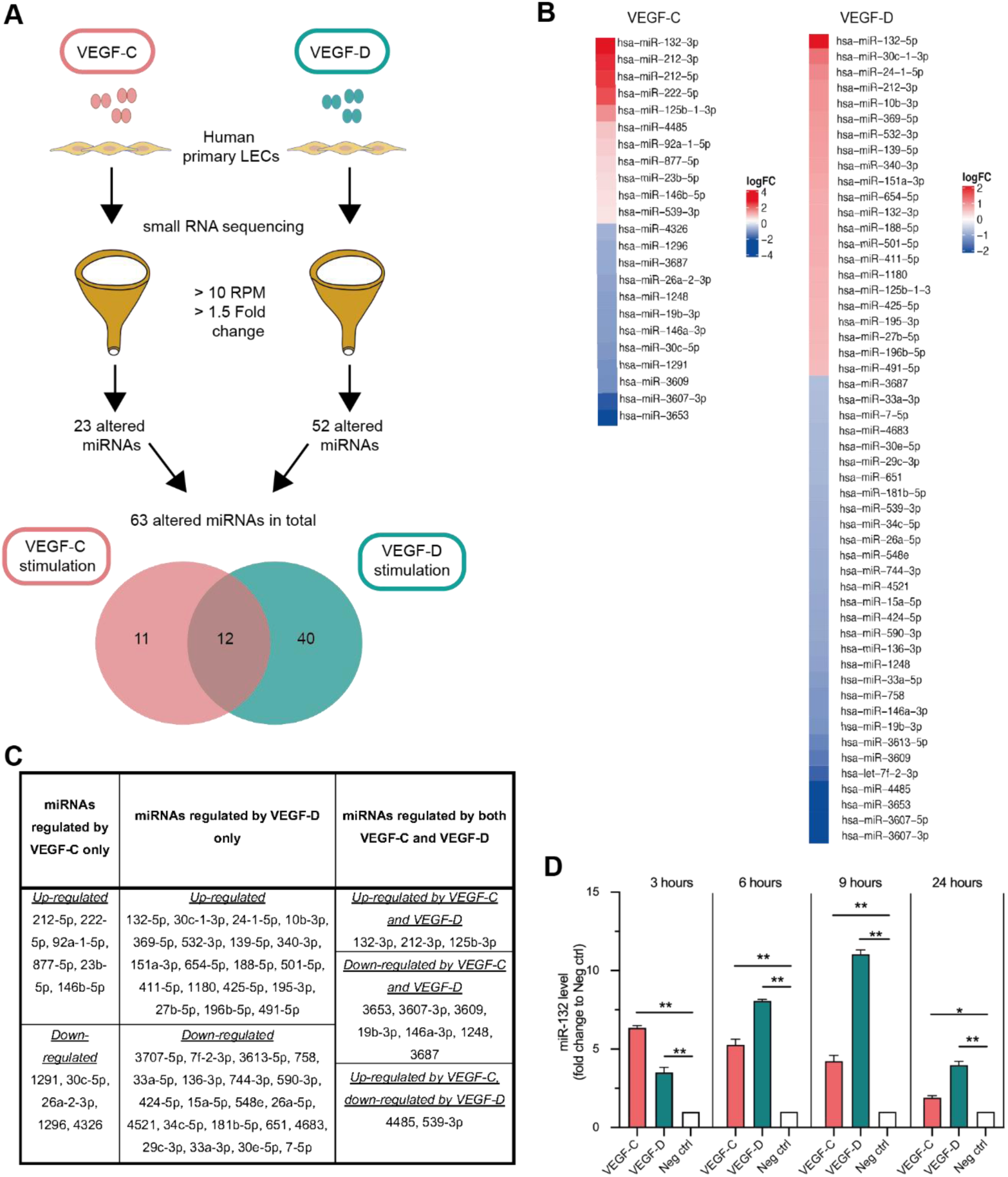
miRNAs regulated by VEGF-C and VEGF-D in LECs. **(A)** Diagram showing the experimental strategy. RPM denotes reads per million. **(B)** Heatmaps showing differential expression of miRNAs in response to VEGF-C or VEGF-D compared to vehicle control. Heatmaps were generated using data from small RNA-Seq and unsupervised hierarchical clustering. **(C)** Table showing miRNAs regulated by VEGF-C, VEGF-D or both. **(D)** RT-qPCR analysis of miR-132 at 3, 6, 9 and 24 h of stimulation of LECs with 200 ng/ml of VEGF-C or VEGF-D. Levels of miR-132 were normalised to vehicle control (“Neg ctrl”; valued at 1.0). Data are mean±SEM from two independent experiments. *P<0.05, ** P<0.01, One-way Anova with Tukey’s post-hoc test.

Before committing to functional studies of miR-132 in lymphangiogenesis, the results of the small RNA-Seq were validated by treating LECs *in vitro* with VEGF-C or VEGF-D for 3, 6, 9 or 24 h, using the same growth factor concentrations and experimental settings as for the RNA-Seq experiment. miR-132 levels were monitored by RT-qPCR which demonstrated a considerable and statistically significant up-regulation of miR-132 in response to VEGF-C compared with untreated LECs at every time-point, with a maximum increase at 3 h of treatment (6.4-fold increase) (Figure 1D). Similarly, levels of miR-132 were increased in response to VEGF-D at every time-point, but with a maximum increase at 9 h of treatment (11-fold increase compared with the negative control). These results confirm the findings from small RNA-Seq that both VEGF-C and VEGF-D induce increased levels of miR-132 in LECs, and demonstrate that the kinetics of these responses are different with peak response being reached faster for VEGF-C than for VEGF-D.

### miR-132 is critical for key steps of lymphangiogenesis *in vitro*

The previous analyses showed that miR-132 is up-regulated in LECs *in vitro* in response to the lymphangiogenic growth factors VEGF-C and VEGF-D. Lymphangiogenesis is a complex, multi-step process that involves proliferation, migration and tube formation by LECs to form new lymphatic vessels (30–32). These distinct steps of lymphangiogenesis can be modelled with *in vitro* assays involving LECs (33). To test the functional role of miR-132 using such assays we employed an antagomiR of miR-132 (designated miR-132 antagomiR #1), delivered to LECs by liposome transfection, to inhibit miR-132. This antagomiR was functionally validated by transfection into LECs followed by monitoring of the mRNA for Retinoblastoma protein (Rb1), a known target of miR-132 (34) by RT-qPCR. Transfection of LECs with miR-132 antagomiR #1, at of concentration of 20 nM, led to a statistically significant increase of Rb1 mRNA to 1.25-fold±0.1-fold (mean±SEM) compared with the Rb1 mRNA level after transfection with a scrambled negative control antagomiR (P<0.05, Student’s t-test), as assessed 48 h post-transfection.

For cell proliferation assays, human primary LECs were transfected with miR-132 antagomiR #1, then stimulated with VEGF-C or VEGF-D, and proliferation was monitored 48 h later. As expected, VEGF-C and VEGF-D both stimulated LEC proliferation (Figure 2A&B). miR-132 antagomiR #1 completely blocked the proliferation of LECs induced by both VEGF-C and VEGF-D whereas a scrambled negative control antagomiR did not (Figure 2A&B). These results indicate that miR-132 is required for LECs to proliferate in response to either VEGF-C or VEGF-D.

**Figure 2.**
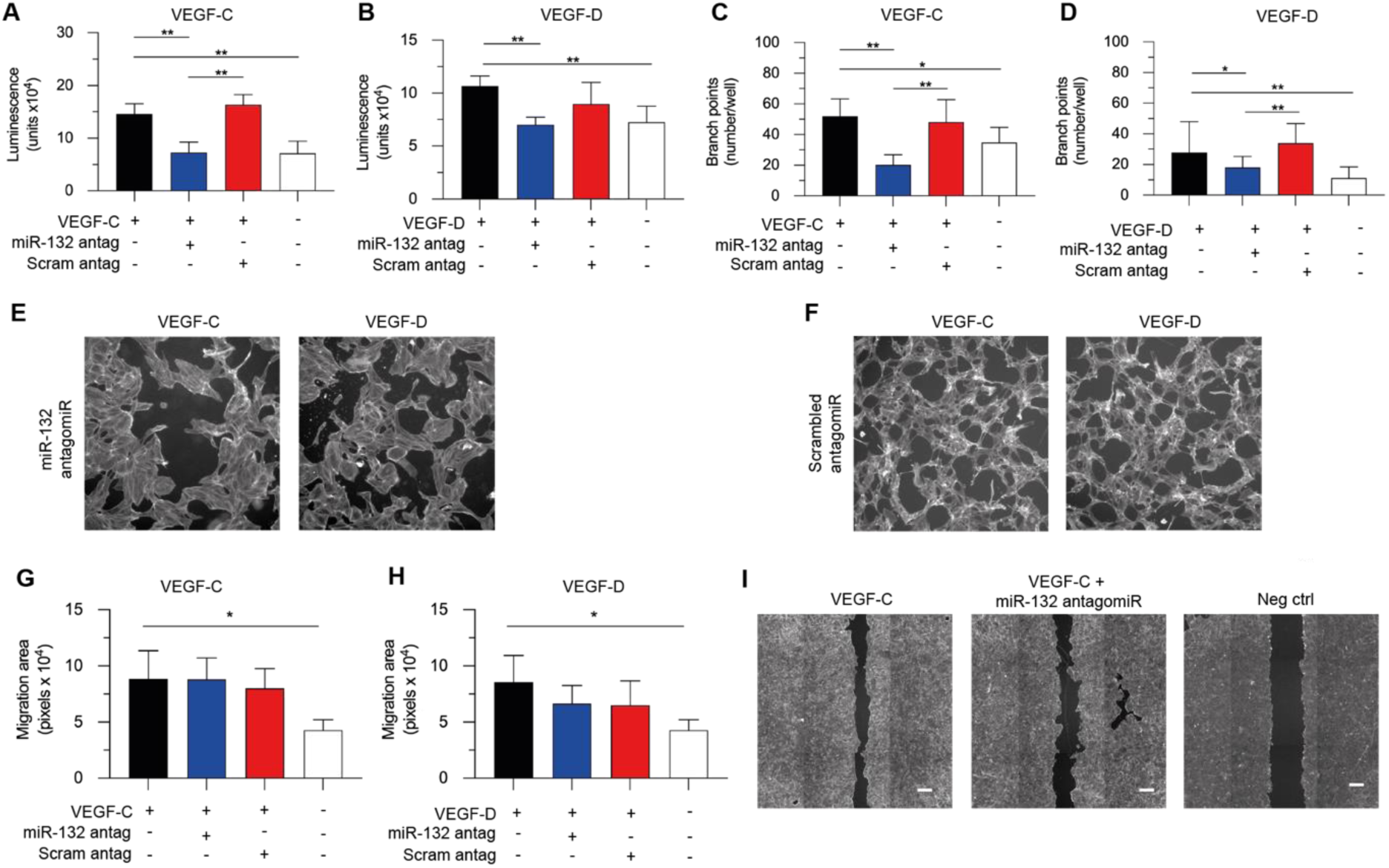
miR-132 regulates key steps of lymphangiogenesis *in vitro*. LECs were transfected with miR-132 antagomiR #1 (“miR-132 antag”) or a scrambled negative control (“Scram antag”) and stimulated with 200 ng/ml VEGF-C **(A)** or VEGF-D **(B)**, and proliferation was measured after 48 h by monitoring luminescence (after addition of a luciferin solution) which is proportionate to the number of viable cells. For analysis of tube formation, LECs were transfected and stimulated as above with VEGF-C **(C)** or VEGF-D (**D**) and tubes monitored after 8 h in a collagen overlay assay by measuring number of tube branch points. **(E,F)** Representative images of tube formation. For analysis of migration, LEC monolayers were transfected as above, scratch wounding was performed, and LECs stimulated with VEGF-C **(G)** or VEGF-D **(H)**, and migration measured after 24 h. **(I)** Representative images of migration assays. Scale bars denote 50 μm. For graphs, data are mean±SEM calculated from three independent experiments. *denotes P<0.05, **P<0.01, One-way Anova with Tukey’s post-hoc test.

The effect of inhibiting miR-132 in LECs on VEGF-C- and VEGF-D-driven formation of vessel-like structures *in vitro* was also monitored. LECs were transfected with miR-132 antagomiR #1, or the scrambled negative control antagomiR, and used in a collagen overlay tube formation assay (16). VEGF-C and VEGF-D both induced tube formation of LECs as assessed by an increase in the number of branch-points of the tubes (Figure 2C&D). Importantly, the inhibition of miR-132 with miR-132 antagomiR #1 significantly restricted branching of the tubes induced by VEGF-C (Figure 2C,E&F) or VEGF-D (Figure 2D,E&F), whereas the scrambled antagomiR control did not.

Finally, the effect of inhibiting miR-132 on LEC migration was monitored in an *in vitro* “wound closure” assay in which a confluent layer of LECs is “scratched” leaving a region devoid of cells into which the LECs can then migrate (35). VEGF-C and VEGF-D both promoted migration of LECs in this assay system (Figure 2G&H). To test the miR-132 inhibitor, LEC monolayers were transfected with miR-132 antagomiR #1, or a scrambled negative control antagomiR, scratched and then incubated with VEGF-C and VEGF-D. Quantification of migrated LECs demonstrated that the miR-132 antagomiR did not significantly alter migration of LECs promoted by VEGF-C or VEGF-D (Figure 2G-I).

Overall, these data demonstrate that miR-132 is required for LEC proliferation and tube formation *in vitro* in response to VEGF-C or VEGF-D but is not required for the migration of LECs induced by either of these growth factors. Therefore, this miRNA is crucial for some, but not all, cell biological aspects of lymphangiogenesis.

### miR-132 is expressed in lymphatics *in vivo*

Based on the findings described above, we hypothesised that miR-132 levels would increase in the LECs of lymphatic vessels in a lymphangiogenic environment *in vivo*, particularly when the lymphatic remodelling is driven by VEGF-C or VEGF-D. We therefore looked for expression of this miRNA in two different pathological models of lymphangiogenesis *in vivo*: (i) Tumour xenografts in mice based on the 293EBNA-1 human embryonic kidney cell line which had been genetically engineered to express full-length human VEGF-D (designated “VEGF-D-293 tumours”) (36); (ii) Human tumours expressing high levels of VEGF-C and/or VEGF-D.

The expression of miR-132 in three VEGF-D-293 tumours was monitored by *in situ* hybridization (ISH), and lymphatics were identified on serial sections by immunostaining for podoplanin, an extensively utilized lymphatic marker broadly expressed on small and large lymphatics (37) (Figure 3A). This revealed that approximately 70% of podoplanin-positive lymphatic vessels detected in these VEGF-D-293 tumours expressed miR-132. More specifically, 20 out of 25 (80%) of podoplanin-positive lymphatics were also positive for miR-132 in the first VEGF-D-293 tumour, 7 out of 12 (58%) were miR-132-positive in the second and 12 out of 17 (71%) were miR-132-positive in the third.

**Figure 3.**
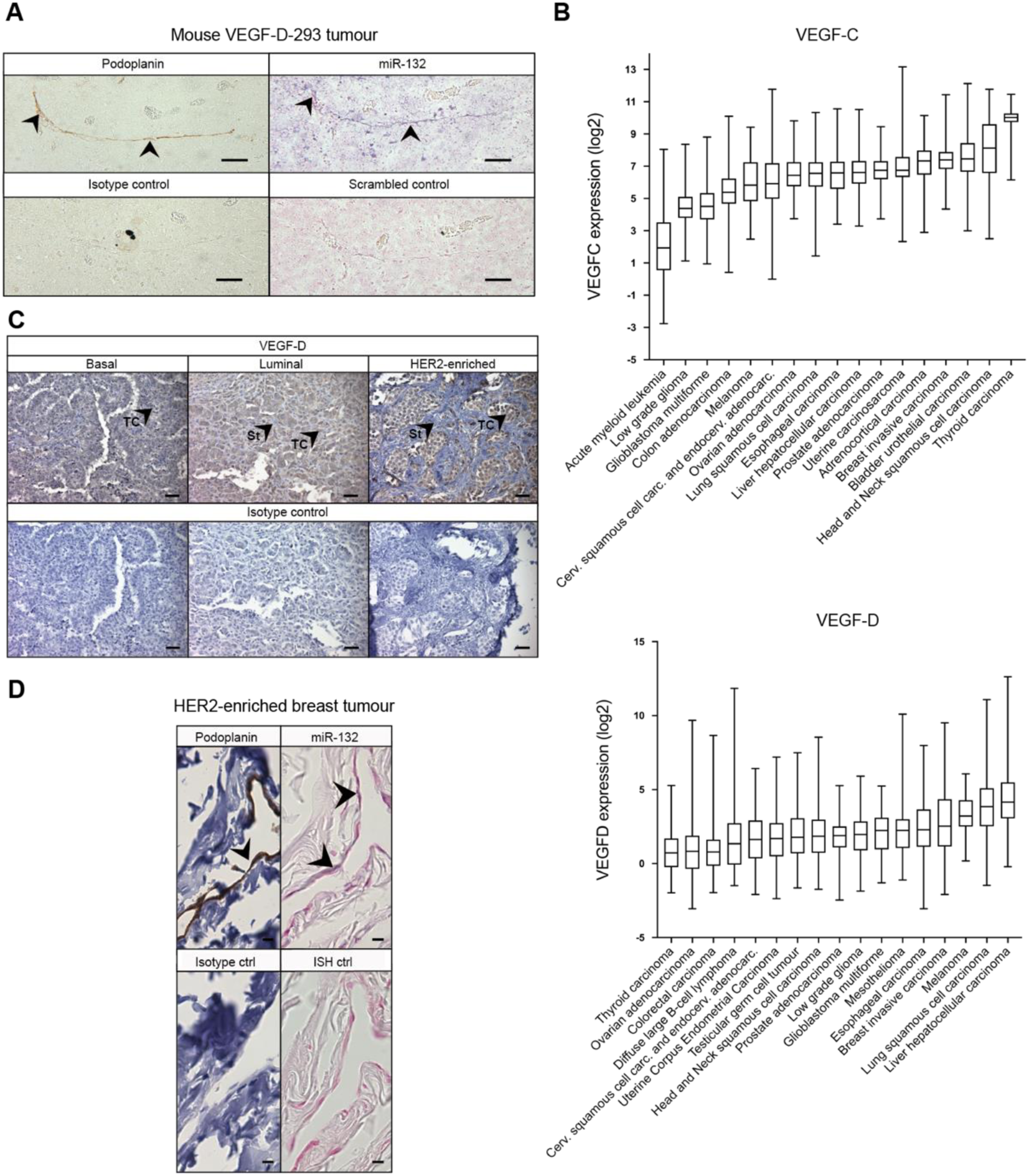
miR-132 is expressed in lymphatics *in vivo*. **(A)** Staining of a lymphatic vessel in a mouse VEGF-D-293 tumour for the lymphatic marker podoplanin (brown) by immunohistochemistry (IHC) and, in serial section, for miR-132 (purple) by *in situ* hybridisation (ISH). Arrowheads: lymphatics; scale bar: 50 μm. Control for podoplanin was isotype-matched antibody and for miR-132 was probe of scrambled sequence, on serial sections. Staining for miR-132 on tumour cells is apparent because VEGF-D-293 cells express this miRNA as determined by RT-qPCR (data not shown). **(B)** mRNA levels for VEGF-C and VEGF-D in 17 human tumour types determined from The Cancer Genome Atlas (TCGA) database (data from TCGA Pan-Can Atlas Study 2018). Data sorted from tumour types with lowest to highest median expression, from left to right. Bars indicate maxima and minima, tops and bottoms of boxes denote upper and lower quartiles, respectively, and lines in boxes denote medians. **(C)** VEGF-D staining (brown) on various types of breast tumours. TC denotes tumour cells and St stroma. Scale bar is 50 μm. Control was isotype-matched antibody, on serial sections. **(D)** Staining of lymphatics for podoplanin (brown/black) by IHC and, in serial section, of miR-132 (purple) by ISH, in a HER2-enriched human breast tumour. Scale bar: 10 μm. Controls as for A. Arrowheads: positive signal on lymphatic endothelium.

Given that VEGF-C and VEGF-D promote expression of miR-132 in LECs, the human tumours likely to contain lymphatic vessels expressing miR-132 would be those in which tumour cells or stromal components secrete VEGF-C and/or VEGF-D. Bioinformatic and immunohistochemical analyses were therefore performed to detect expression of these growth factors in a variety of human tumour types. Bioinformatic analysis was performed by examining the publicly available gene expression database for human tumours, The Cancer Genome Atlas (TCGA). The RNA sequencing data from TCGA was used to assess VEGF-C and VEGF-D mRNA levels in 17 tumour types (Figure 3B). This analysis revealed that on average, the levels of VEGF-C mRNA are higher in thyroid carcinoma, head and neck squamous cell carcinoma, bladder urothelial carcinoma and breast invasive carcinoma compared with the other tumour types analysed, whereas VEGF-D mRNA is more abundant in liver hepatocellular carcinoma, lung squamous cell carcinoma, melanoma and breast invasive carcinoma. Breast invasive carcinoma was chosen for immunohistochemical analysis because both VEGF-C and VEGF-D mRNAs are commonly detected at relatively high levels in these tumours. Subsequent to bioinformatic analysis, immunohistochemical analysis of breast cancer tissue microarrays (TMAs) containing human breast invasive carcinomas was performed to identify individual tumours expressing VEGF-D; immunohistochemistry for VEGF-C was not performed due to the lack of available antibodies which, in our hands, could reliably detect this protein in human tissues. The breast cancer TMAs contained 120 different tumour samples which included different subtypes of invasive breast cancer, specifically luminal A and B breast cancer, basal-like breast cancer and human epidermal growth factor receptor 2 (HER2)-amplified breast cancer. The analysis of VEGF-D by immunohistochemistry (for example see Figure 3C) demonstrated that approximately 80% of HER2-enriched breast cancers showed high intensity of VEGF-D staining which was by far the highest percentage for any of the tumour types analysed. These data suggested that VEGF-D-positive HER2-enriched breast cancers would be an appropriate choice for analysing miR-132 in tumour lymphatics. Therefore, two HER-2-enriched breast cancers expressing high levels of VEGF-D were chosen for analysis of miR-132. Tissue sections of these tumours were stained by immunohistochemistry for podoplanin to highlight lymphatic vessels, and serial sections were subjected to ISH for miR-132 (Figure 3D). Approximately 19% (4 out of 21) of podoplanin-positive lymphatic vessels detected in one of these HER2-enriched tumours were also positive for miR-132, whereas none of 20 podoplanin-positive lymphatics detected in the second tumour were positive for miR-132.

These data demonstrate that solid tumours in mice and humans, expressing VEGF-D, can contain lymphatic vessels which are positive for miR-132. The heterogeneity of miR-132 levels in tumour lymphatics which we observed could be due to variation in VEGF-D concentrations in different parts of a tumour. Further, VEGF-D requires proteolytic activation to bind VEGFR-3 on LECs (38) so its capacity to drive expression of miR-132 in tumour lymphatics is dependent on the proteases which activate it (39, 40) – the levels of these proteases may be highly variable within and between tumours.

### miR-132 regulates lymphangiogenesis *in vivo*

In order to assess if miR-132 regulates lymphangiogenesis *in vivo*, we employed a mouse ear model of lymphangiogenesis which has been shown previously to be robust, reproducible and to allow extensive morphological characterisation of the lymphatic network (17, 41) (Figure 4A). To test the effects of inhibiting miR-132 in this model, a miR-132 antagomiR was employed (designated miR-132 antagomiR #2) which was bound to a cholesterol molecule to facilitate entry of the antagomiR into cells *in vivo*. Before using this antagomiR *in vivo*, its efficacy was tested *in vitro* in LECs at four different concentrations, and it was shown to significantly up-regulate mRNA levels for a validated target of miR-132, Rb1, when used for transfection of LECs at a concentration of 40 nM (Figure 4B).

**Figure 4.**
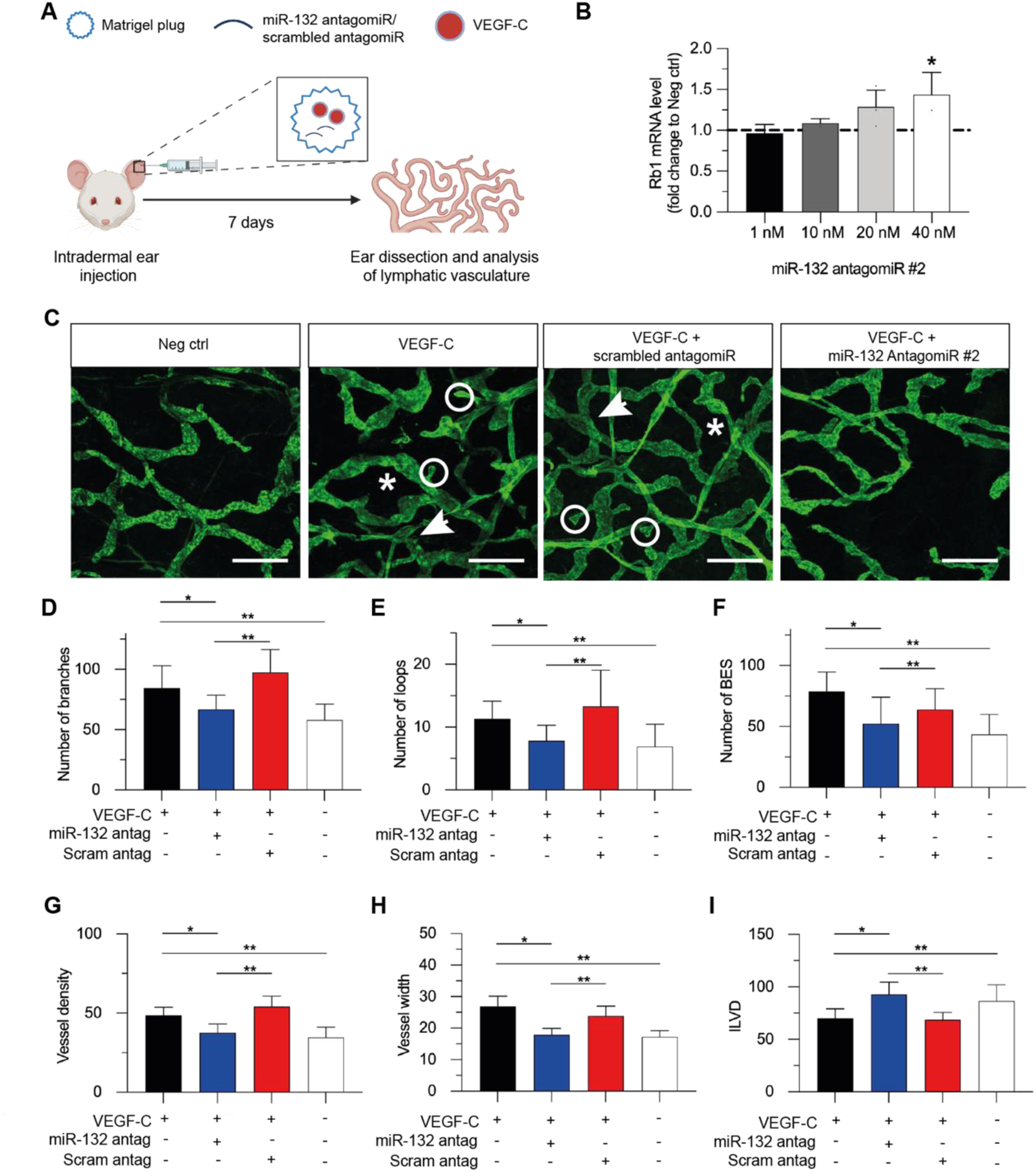
miR-132 regulates lymphangiogenesis *in vivo* in a mouse ear model. **(A)** Schematic representation of the experiment. **(B)** Validation of antagomiR. mRNA for Rb1, a known target of miR-132, was monitored by RT-qPCR 48 h post-transfection of LECs with miR-132 antagomiR #2, or scrambled negative control antagomiR, at four concentrations. Results for negative control indicated by slashed line. Data are mean±SEM from two independent experiments. *P<0.05, Student’s t-test. **(C)** Representative confocal images of ears, stained for LYVE1, for the four study groups. Asterisks denote lymphatic loops; circles, blind-ended sacs; arrows, branch-points; scale bar, 50 µm. Parameters for quantifying lymphatic morphology were vessel branches **(D)**, loops **(E)**, blind-ended sacs (“BES”) **(F)**, vessel density **(G)**, vessel width **(H)** and inter-lymphatic vessel distance (“ILVD”) **(I)**. Numbers in D-G indicate branches, loops, BES and vessels, respectively, per 4 high-powered fields per ear, and in H and I indicate width and distance, respectively, in pixels. “miR-132 antag” denotes miR-132 antagomiR #2; “Scram antag”, scrambled antagomir. Quantification involved 16 to 19 ears per study group across two independent experiments. Data are mean±SEM. *P<0.05, **P<0.01, One-way Anova with Tukey’s post-hoc test.

The mouse ear lymphangiogenesis assay involved intradermal injection of recombinant, mature human VEGF-C, mixed with Matrigel, into the ears of SCID/NOD/gamma (NSG) mice to establish a Matrigel plug which stimulated lymphangiogenesis in the surrounding tissue. In order to test whether inhibiting the action of miR-132 blocked VEGF-C-induced lymphangiogenesis in this assay, mice were injected with Matrigel and VEGF-C, in combination with miR-132 antagomiR #2 or a scrambled negative control antagomir. Matrigel co-injected with VEGF-C only was the positive control and Matrigel co-injected with vehicle only was the negative control. Mice were sacrificed one week after injections and their ears were excised, fixed and dissected to expose the dermal layer of skin and subsequently perform whole-mount immunostaining for the lymphatic vessel marker LYVE-1. LYVE-1 staining revealed that targeting miR-132 with miR-132 antagomiR #2 significantly restricted VEGF-C-induced lymphangiogenesis and lymphatic remodelling in the ear (Figure 4C). A range of morphological characteristics of the lymphatic network was monitored using specialised software developed for analysis of the lymphatic vasculature (42). Specifically, VEGF-C increased the number of lymphatic branches (Figure 4D), loops (Figure 4E) and blind-ended sacs (Figure 4F). VEGF-C also increased lymphatic vessel density (Figure 4G) and width (Figure 4H), and reduced the distance between lymphatic vessels (Figure 4I). In contrast, when VEGF-C was co-injected with miR-132 antagomiR #2, there was a statistically significant reduction in all of these parameters compared with VEGF-C treatment alone or co-treatment with VEGF-C and the negative control antagomiR. In summary, targeting of miR-132 inhibited many aspects of lymphatic remodelling induced by VEGF-C in the mouse ear lymphangiogenesis assay.

### Identification of miR-132 interactome in LECs

To understand how miR-132 mechanistically controls the molecular pathways involved in lymphangiogenesis, we identified the interactome of this miRNA in LECs using Argonaute High Throughput Sequencing after Cross-Linked Immunoprecipitation (Ago HITS-CLIP) technology. This approach allows the global identification of mRNAs regulated by a miRNA in a cell. After transfection with a molecular mimic of the miRNA, RNA-protein complexes in the cell are cross-linked, Ago (the protein on which miRNAs pair with mRNA targets) is immunoprecipitated along with mRNAs bound to it, and these mRNAs are isolated for sequencing to identify mRNAs which are enriched due to the presence of the mimic (Figure 5A) (43). We transfected human primary LECs with a miR-132 mimic (a chemically modified double-stranded RNA molecule that mimics endogenous miR-132), or a scrambled mimic negative control, and then performed Ago HITS-CLIP. From this experiment, we identified approximately 500 mRNAs putatively regulated by miR-132 as CLIP peaks enriched in miR-132-transfected cells compared to cells transfected with the negative control (Supplementary Datafile 1 and Supplementary Figure 1). Some mRNA targets identified by the Ago HITS-CLIP were previously validated targets of miR-132, for instance mRNAs for the phosphatase PTEN (44) and the cyclin-dependent kinase inhibitor p21 (45). These targets validated the experiment and provided some immediate insight into mRNAs regulated by miR-132 in LECs.

**Figure 5.**
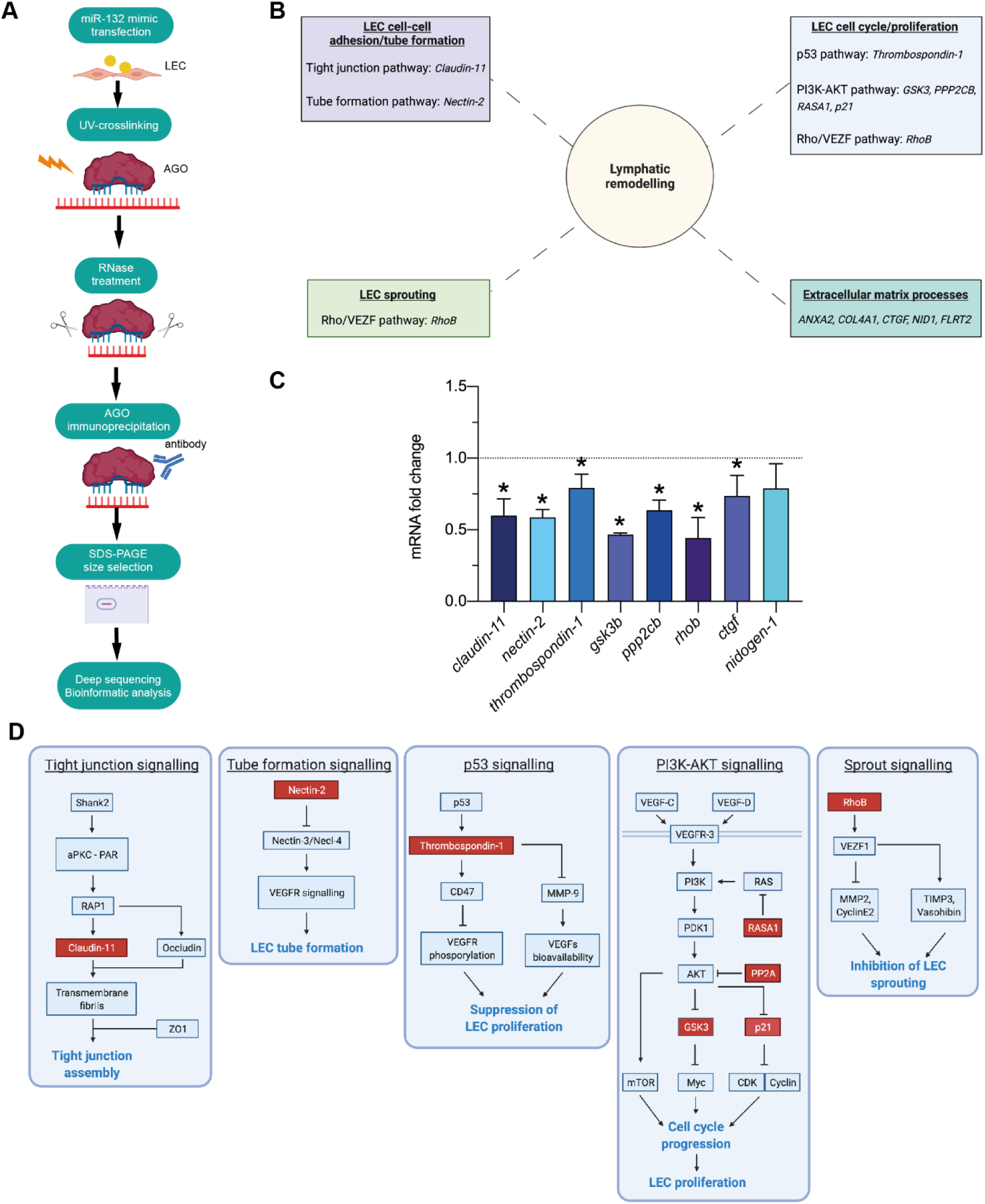
Identification of mRNAs regulated by miR-132 in LECs by Ago HITS-CLIP. **(A)** Schematic of Ago HITS-CLIP. miR-132 mimic combined with liposomes: yellow circles; miRNA mimic: blue; mRNA: red. **(B)** Pathways and genes, regulated by miR-132 and involved in lymphatic remodelling, as identified by Ago-HITS-CLIP and bioinfomatics. Genes are indicated by italic text. **(C)** LECs were transfected with miR-132 mimic, and levels of mRNAs assessed by RT-qPCR. Dotted line: results with scrambled negative control mimic. Data are mean±SEM. *P<0.05, Student’s t-test, three independent experiments. “ppp2cb” denotes mRNA for PPP2CB which is the beta isoform of the catalytic subunit of the PP2A phosphatase. “ctgf” denotes mRNA for connective tissue growth factor. **(D)** Proteins for which their encoding mRNAs are regulated by miR-132 in LECs (red) in signalling pathways relevant to lymphatic remodelling (adapted from Kyoto Encyclopedia of Genes and Genomes). For PP2A in the PI3K-AKT signalling diagram, it is the mRNA for its catalytic subunit, PPP2CB, that is regulated by miR-132.

**Supplementary Figure 1.**
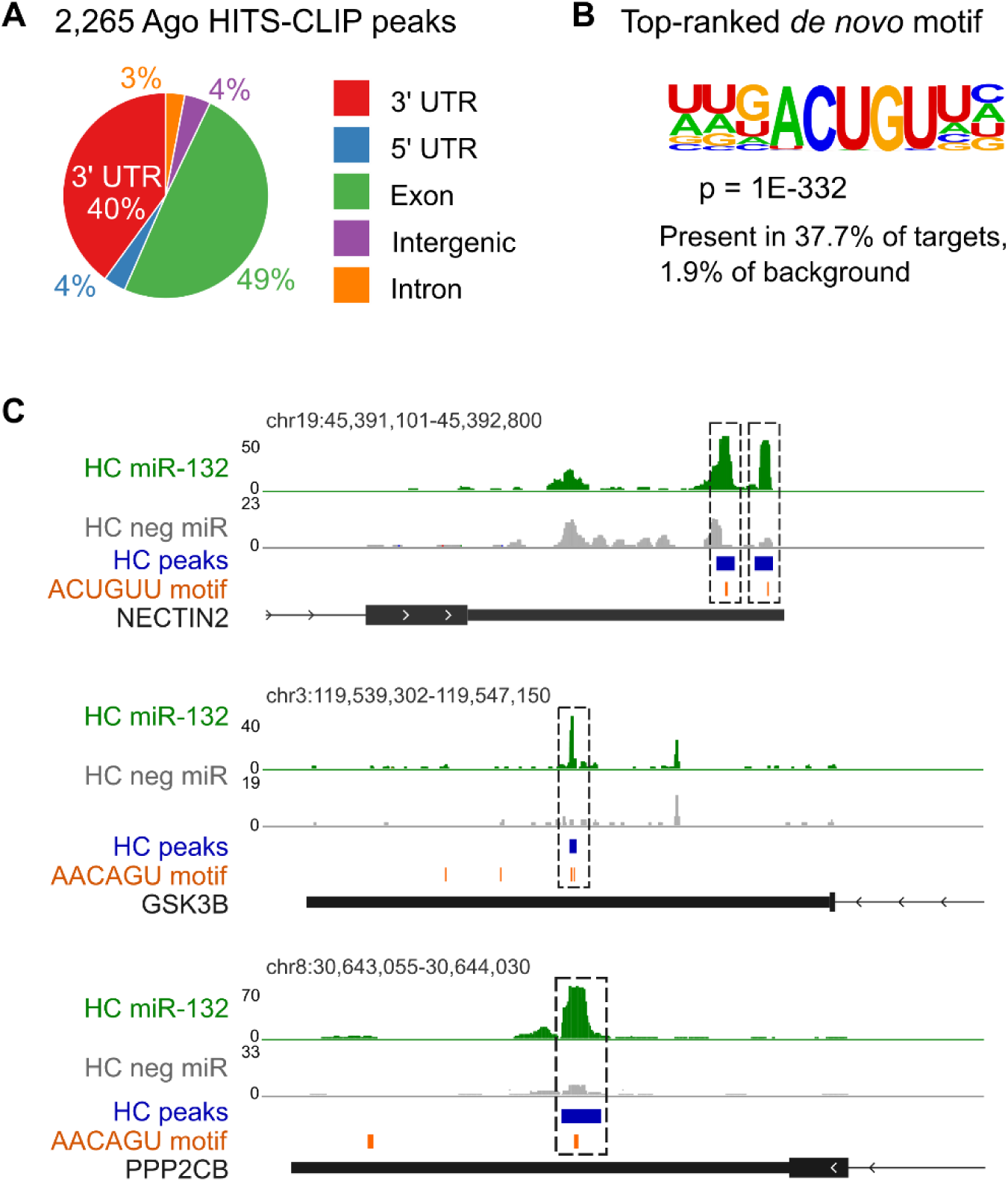
HITS-CLIP read alignments from samples transfected with miR-132 mimic form enriched peaks at potential miR-132 binding sites. **(A)** Distribution of 2,265 Ago-HITS-CLIP peaks mapped to their genomic regions. “UTR” denotes untranslated. **(B)** Top-ranked motif identified from *de novo* motif discovery from peaks in 3’ UTR regions. **(C)** Examples of HITS-CLIP read alignment peaks in the 3’ UTR regions of NECTIN2, GSK3B and PPP2CB. Peaks of reads enriched in LECs transfected with miR-132 mimic (indicated by “HC miR-132” in green) relative to those transfected with a scrambled mimic negative control (indicated by “HC neg miR” in grey) and motifs complementary to the miR-132 core seed region (positions 2-7) are shown in blue and orange, respectively. Coverage track limits are scaled to normalise for sequencing depth.

In order to analyse the results from the Ago HITS-CLIP, we performed pathway and gene ontology analyses using the Database for Annotation, Visualization and Integrated Discovery (DAVID), an online tool to identify enriched biological pathways from gene lists (https://david.ncifcrf.gov/) (46), and Gene Ontology (GO), a database on gene function (http://geneontology.org/) (47), respectively. The pathway analysis was conducted on all genes from the Ago HITS-CLIP, and demonstrated enrichment of molecular pathways relating to cell cycle, cell-cell junctions, mRNA processing and protein ubiquitination. Genes and pathways for further consideration were then chosen by performing gene ontology analysis thus identifying pathways, from those highlighted by pathway analysis, which were most likely to be involved in lymphangiogenesis and lymphatic remodelling – these pathways are listed in Figure 5B. To evaluate the reliability of our HITS-CLIP results, we transfected LECs with the miR-132 mimic and checked by RT-qPCR whether eight of the mRNAs, randomly chosen from mRNAs identified by Ago HITS-CLIP and predicted by our bioinformatic analyses to be involved in lymphagiogenesis, were down-regulated. These mRNAs were for Claudin-11, Nectin-2, Thrombospondin-1, GSK3B (glycogen synthase kinase 3 beta), PPP2CB (the beta isoform of the catalytic subunit of protein phosphatase 2 (PP2A)), RHOB, CTGF (connective tissue growth factor) and Nidogen-1. Figure 5C shows that the levels of all eight mRNA targets were down-regulated by the miR-132 mimic, as expected, with the reduction being statistically significant for seven, namely the mRNAs for Claudin-11, Nectin-2, Thrombospondin-1, GSK3B, PPP2CB, RHOB and CTGF. These results indicate a high degree of reliability for the Ago HITS-CLIP experiment.

The analyses described above identified multiple pathways via which miR-132 co-ordinately regulates key steps in lymphangiogenesis and lymphatic remodelling, including LEC proliferation and tube formation (Figure 5D). The down-regulation by miR-132 of mRNA for Claudin-11 in LECs (Figure 5C) is of particular interest because this protein is an important component of tight junctions (48), which are prominent in the junctions between LECs in lymphatic vessels (49) and play a key role in mediating LEC cell-cell adhesion (50). Down-regulation of Claudin-11 would likely restrict formation of tight junctions and favour remodelling of LECs to generate tube-like structures, as was reported to analogously occur when Claudin-5 was down-regulated in blood vascular endothelial cells (51). Nectin-2 has been reported to be a negative regulator of tube formation by endothelial cells (52) (Figure 5D) so down-regulation of its expression by miR-132 (Figure 5C), when LECs are exposed to VEGF-C or VEGF-D, would also favour LEC tube formation. Thrombospondin-1 is an inhibitor of lymphangiogenesis (53) which can act by restricting endothelial cell proliferation (54) (Figure 5D). Thus, the down-regulation of mRNA for Thromobospondin-1 in LECs by miR-132 (Figure 5C) would favour LEC proliferation. It is noteworthy that four of the miR-132-regulated mRNAs identified by Ago HITS-CLIP encode GSK3B, PPP2CB (a catalytic subunit of PP2A), RASA1 (protein p120 Ras GTPase-activating protein) and P21 which are all associated with the PI3K-AKT signalling pathway (Figure 5D). This pathway, like miR-132, is positively regulated in LECs by VEGF-C and VEGF-D, and is important for proliferation of LECs and lymphangiogenesis (15, 55). All four proteins are negative regulators of the PI3K-AKT pathway, indicating that the up-regulation of miR-132 in LECs by VEGF-C or VEGF-D, leading to down-regulated expression of these proteins, would co-ordinately enhance activity of this key lymphangiogenic signalling pathway via four distinct control points (Figure 5D). Finally, RHOB is a negative regulator of lymphangiogenesis which acts by restricting LEC sprout formation (56) (Figure 5D). The mRNA for RHOB is down-regulated in LECs by miR-132 (Figure 5C), so higher levels of miR-132 in LECs driven by VEGF-C or VEGF-D would favour lymphatic sprouting which is another important step in lymphatic remodelling.

## DISCUSSION

We used an approach based on small RNA-Seq to identify miRNAs regulated by the lymphangiogenic growth factors VEGF-C and VEGF-D, with a view to better understanding how signalling for lymphatic remodelling is co-ordinately regulated in LECs. This revealed that VEGF-C and VEGF-D significantly altered the expression of 63 miRNAs in LECs, with some miRNAs being up-regulated and others down-regulated. Interestingly, only 12 of those were regulated by both growth factors, indicating that VEGF-C and VEGF-D can exhibit distinct signalling in LECs despite their structural homology and similar receptor binding properties. This distinct signalling was previously observed by Karnezis and co-workers who found that VEGF-C and VEGF-D were able to stimulate different signalling pathways in lymphatic vessels (57, 58). Both VEGF-C and VEGF-D can bind VEGFR-2, as well as VEGFR-3, on LECs, and it has been reported that they can promote formation of VEGFR-2 homodimers, VEGFR-3 homodimers and VEGFR-2/VEGFR-3 heterodimers (59–63). Each of these receptor complexes could potentially induce distinct patterns of signalling (61) leading to regulation of different groups of miRNAs. Thus, if VEGF-C and VEGF-D have different propensities for generating each receptor homodimer, or the receptor heterodimer, this could explain how these growth factors regulate distinct, but partly overlapping, groups of miRNAs. It has also been reported that VEGF-C and VEGF-D can bind non-VEGF receptors, and promote formation of VEGF receptor complexes involving co-receptors, including neuropilins and integrins (64–67). If VEGF-C and VEGF-D have different propensities for binding such non-VEGFR cell surface molecules, this might also contribute to distinct patterns of signalling leading to regulation of different groups of miRNAs. The molecular mechanisms by which VEGF-C and VEGF-D signalling regulate levels of miRNAs in LECs, which could involve modulation of miRNA biosynthesis or degradation, are unknown and warrant investigation.

We showed that miR-132 was the most up-regulated miR in LECs in response to three hours of exposure to VEGF-C, and was significantly up-regulated by VEGF-D over the same time period. However, the kinetics of the responses to these proteins differed in that the peak in miR-132 level in response to VEGF-C occurred much sooner than for VEGF-D. The reason for the different kinetics of these responses is unknown but could relate to different affinities of VEGF-C and VEGF-D for VEGF receptor homodimers versus heterodimers, and different affinities for cell surface non-VEGFR molecules such as neuropilins, integrins and heparan sulphate proteoglycans. miR-132 was critical for VEGF-C- and VEGF-D-driven proliferation and tube formation by LECs *in vitro*, but not for migration of these cells. This suggests that the VEGF-C- and VEGF-D-driven pathways signalling for LEC migration are distinct from those controlling proliferation and tube formation, and that there is a lack of miR-132 targets in the migratory pathways. Targeting miR-132 *in vivo* in mice blocked many morphological features of VEGF-C-driven lymphatic remodelling, including vessel branching, and increases in vessel width and density, likely reflecting the involvement of LEC proliferation and tube formation in multiple aspects of lymphatic vessel remodelling. miR-132 has also been found to stimulate remodelling of blood vessels *in vivo* in a mouse model of breast carcinoma (28). Taken together, these results indicate that miR-132 is a “master regulator” of remodelling across the blood and lymphatic vasculatures, likely exploiting similar molecular mechanisms in both settings.

In addition to unravelling the crucial function of miR-132 in lymphatic remodelling, our study identified mechanisms by which this miRNA regulates this complex process. The miR-132-mRNA interactome was identified in LECs by Ago HITS-CLIP indicating that miR-132 controls multiple molecular pathways critical for VEGF-C- or VEGF-D-driven lymphatic remodelling, including pathways important for the biological processes of LEC proliferation and tube formation, as well as lymphatic branching which is dependent on sprout formation; these processes were shown to be dependent on miR-132 in our *in vitro* and *in vivo* studies (Figures 2A-D and 4D). More specifically, a miR-132 mimic down-regulated the mRNAs for Claudin-11 and Nectin-2 in LECs which would co-ordinately modulate two distinct signalling pathways to favour tube formation (51, 52) (Figure 5D). Likewise, the miR-132 mimic down-regulated, in LECs, the mRNAs for Thrombospondin-1 and proteins that negatively regulate the PI3K-AKT pathway, which would co-ordinately modulate two signalling pathways to favour LEC proliferation (15, 53) (Figure 5D). The down-regulation we observed of the mRNA for RHOB in LECs, in response to the miR-132 mimic, would favour formation of lymphatic sprouts (56), another key step in lymphatic remodelling. These findings indicate that miR-132 can (i) co-ordinately modulate an important signalling pathway for lymphatic remodelling at multiple control points (e.g. the PI3K-AKT pathway for LEC proliferation); (ii) co-ordinately modulate multiple signalling pathways to promote a biological process in lymphatic remodelling (e.g. the PI3K-AKT and Thrombospondin-1 pathways involved in LEC proliferation); (iii) co-ordinately modulate multiple processes in lymphatic remodelling (e.g. LEC proliferation, tube formation and sprouting).

Our finding that miR-132 is a key regulator of lymphangiogenesis could have clinical implications as it may be a useful therapeutic target for diseases in which lymphangiogenesis and lymphatic remodelling play a role in disease progression, such as cancer, lymphoedema and lymphangioleiomyomatosis. miR-132 could potentially be targeted systemically *in vivo* by nucleic acid-based therapeutics. Such therapeutics have faced several hurdles, especially related to the challenges in achieving efficient delivery to organs other than the liver, and overcoming off-target effects and chemistry-dependent toxicity. Nevertheless, up to 2021, 11 oligonucleotide-based therapies had been approved for clinical use (68), and a lipid-nanoparticle formulated, nucleoside-modified synthetic RNA for immunisation against SARS-CoV-2 (the BNT162b2 mRNA Covid-19 vaccine) (69) recently achieved regulatory approval. Also, many more therapeutics based on nucleic acids are being assessed in clinical trials (68). These developments are encouraging for the feasibility of targeting miR-132 in the clinic. As miR-132 has the ability to regulate entire networks of pathways by controlling the expression of many mRNA targets in multiple pathways, it could be an appropriate target for comprehensively inhibiting lymphangiogenesis and lymphatic remodelling. The targeting of a specific growth factor, such as VEGF-C or VEGF-D, or a specific growth factor receptor, such as VEGFR-3, might encounter inherent or acquired drug resistance because alternative lymphangiogenic growth factors or receptors are expressed in a disease prior to treatment, or become up-regulated in response to therapy. In contrast, targeting miR-132 may be less likely to encounter resistance given this miRNA targets multiple lymphangiogenic signalling proteins/pathways which potentially act downstream of a variety of lymphangiogenic growth factors and receptors.

Our findings have highlighted a miRNA which co-ordinates the positive regulation of multiple cell biological events in lymphatic remodelling, and is thus important for lymphangiogenesis *in vivo*. Furthermore, our discovery of other miRNAs regulated by the lymphangiogenic factors VEGF-C and VEGF-D provides a platform for unravelling other mechanisms regulating lymphangiogenesis and lymphatic remodelling, including identification of new signalling pathways critical for these complex biological processes. Moreover, as new signalling pathways for vascular remodelling in disease are discovered from whole-genome functional screens (17, 70), our platform will likely facilitate identification of molecular mechanisms by which these new pathways are co-ordinately regulated to drive vascular remodelling. Finally, our platform may also prove useful for identifying novel therapeutic targets, potentially miRNAs or proteins, for stimulating or restricting lymphatic vessel remodelling in disease.

## MATERIALS AND METHODS

### Cell lines and culture conditions

Cell lines were grown in a humidified atmosphere at 5% CO_2_, or where specified at 8% CO_2_, at 37°C. Human dermal LECs (PromoCell, Heidelberg, Germany), referred to here as “LECs”, are primary cells derived from human foreskin and skin tissues. LECs were cultured on a substrate of human fibronectin (Corning) in complete medium (EGM-2-MV, Lonza, Basel, Switzerland) consisting of basal medium (EBM-2) supplemented with 5% fetal bovine serum (FBS), hydrocortisone, fibroblast growth factor, VEGF, R3 insulin-like growth factor, epidermal growth factor, gentamicin/amphotericin B (GA-1000) and ascorbic acid, at proprietary concentrations. Immortalised mouse mesenteric lymph node endothelial cells (“imLECs”) were cultured in EGM-2-MV in a humidified atmosphere of 8% CO_2_ at 37°C. 293-EBNA-1 human embryonic kidney cells expressing full-length human VEGF-D (“293-EBNA-1-VEGF-D cells”) were established previously (36) and cultured in Dulbecco’s modified Eagle’s medium (DMEM, Thermo Fisher Scientific), 10% FBS, 100 U/ml penicillin and 100 μg/ml streptomycin.

### miRNA antagomiRs and mimics

The mirVana miR-132 antagomiR #1, mimics and negative controls (Invitrogen, Carlsbad, CA) were resuspended in RNase-free H_2_O at 50 μM for *in vitro* cell transfections. miR-132 antagomiR #2 and negative control (GenePharma, Shanghai, China) for *in vivo* use were single-stranded RNAs, coupled to cholesterol, with four thiol modifications at the 3’-terminus and two at the 5’-terminus; the entire strand was modified by 2′-O-methylation. They were resuspended in RNase-free H_2_O at 1 mM.

### Cell transfection and stimulation with growth factors

LECs were transfected with mirVana miRNA antagomiRs, mimics or negative controls (Invitrogen) at 20 nM, using the reverse transfection method as follows. Lipofectamine RNAiMAX was diluted into Opti-MEM according to manufacturer’s instructions (Thermo Fisher Scientific). mirVana miRNA antagomiRs were also diluted in Opti-MEM. These solutions were then combined, incubated for 5 min at room temperature and added to a 96-, 12- or 6-well plate, and overlaid with the appropriate number of cells previously mixed in EGM-2-MV not containing antibiotics. The medium was replaced, 24 h post-transfection, with fresh EGM-2MV for semi-quantitative PCR or with starvation medium if cells were subsequently treated with growth factors.

For treatment with growth factors, LECs and imLECs were starved in serum starvation medium (EBM-2 with hydrocortisone, ascorbic acid, GA-1000 all at proprietary concentrations and 2% bovine serum albumin (BSA, Sigma-Aldrich)) for 24 h, followed by stimulation with serum starvation medium supplemented with recombinant mature human VEGF-C (Opthea, Melbourne, Australia) or recombinant mature human VEGF-D (Opthea) at 200 ng/ml. Stimulation with growth factors was for 3, 6, 9 or 24 h.

### RNA and microRNA isolation

Total RNA was isolated using a silica membrane RNA purification method (RNeasy Mini Kit, Qiagen, Hilden, Germany) according to the manufacturer’s protocol. miRNAs were isolated using the miRNeasy Mini Kit (Qiagen) according to manufacturer’s protocol. The concentration and purity of RNA and miRNA samples were assessed using a nanovolume spectrophotometer (NanoDrop^TM^ 2000, Thermo Fisher Scientific).

### Complementary DNA synthesis and real-time PCR

cDNA synthesis with RNA samples was performed by reverse transcription using random primers (High-Capacity cDNA Reverse Transcription Kit, Thermo Fisher Scientific) according to manufacturer’s protocol. cDNA synthesis with miRNA samples was performed via poly-A tailing and 5’ ligation of an adaptor sequence to extend mature miRNAs on each end prior to reverse transcription. Universal RT primers recognized common sequences present on both the 5’- and 3’-extended ends of the mature miRNAs. Mature miRNAs were reverse transcribed to cDNA (TaqMan™ Advanced miRNA cDNA Synthesis Kit, Applied Biosystems, Waltham, MA, USA) according to manufacturer’s protocol. RT-qPCR was conducted with SYBR green detection (Thermo Fisher Scientific) according to manufacturer’s protocol (primers listed in Table 1) - each sample was run in triplicate reactions in 96-well reaction plates (MicroAmp® Fast Optical, Thermo Fisher Scientific) in a 96-well Real-Time PCR instrument (StepOnePlus^TM^ Real-Time PCR System, Thermo Fisher Scientific). mRNA levels were normalised to mRNA for human β-actin, and miRNA levels were normalised to levels of hsa-miR-423-5p, hsa-miR-345-5p and hsa-let7g-5p. Data were analysed using the comparative C_T_ method as described (71).

**Table 1:**
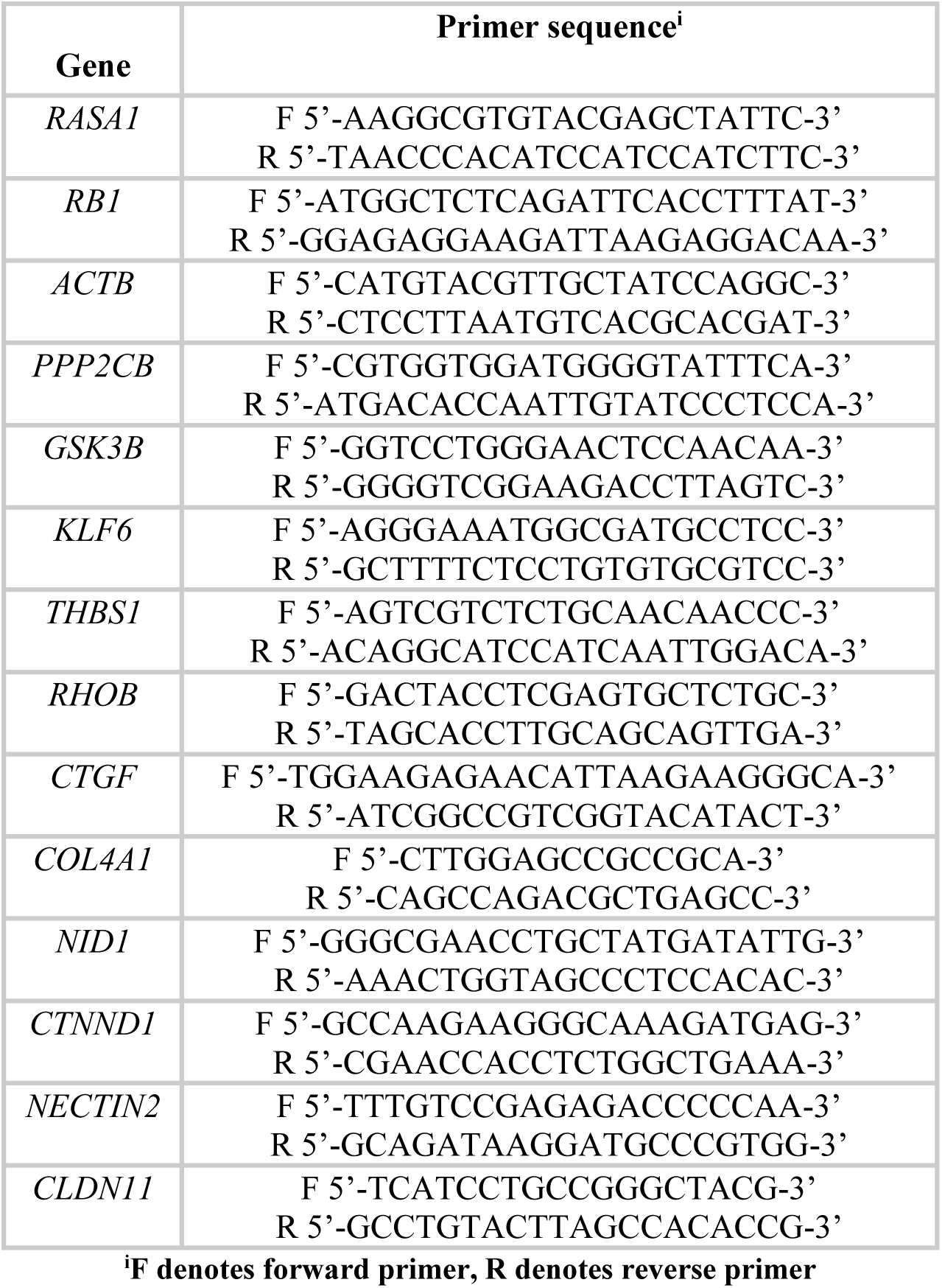
Primers used for reverse transcription real-time PCR.

### Cell proliferation assay

LECs were plated in black-walled 96-well imaging plates (Costar ®, Corning) at 7,000 cells per well, and transfected with miRNA mimics or antagomiRs. After 24 h, cells were washed in PBS and treated with growth factors. At 72 h post-transfection, cell number was assessed using a CellTiter-Glo® Luminescent Cell Viability Assay (Promega, Madison, WI), according to manufacturer’s instructions. Luminescence values were read using a multi-well luminometer (Cytation 3 Cell Imaging Multi•Mode Reader_, B_ioTek Instruments, Winooski, Vermont).

### Cell migration assay

LECs were plated in black-walled 96-well imaging plates (Costar ®, Corning) at 1.5 x 10^4^ cells per well, and transfected with miR mimics or antagomiRs. After 24 h, cells were serum starved and, 48 h post-transfection, a 96-pin wounding device (tool belonging to the Sciclone ALH 3000 Workstation, Caliper Life Sciences, Waltham, MA) was used to create uniform scratches in cell monolayers. Experiments were carried out on duplicate plates, one whose end-point was immediately after the scratch (the *t_0_* plate) and one whose end-point was after 24 h of treatment with growth factors as described below (the *t*_24_ plate). Following the scratch, cells in the *t*_24_ plate were washed once with PBS and then treated with growth factors for 24 h. At the assay end-points, both plates were washed once with PBS and fixed with 1% PFA for 2 h at room temperature. Cells were then washed in PBS, permeabilised and stained using a PBS solution containing 2% BSA, 0.2% Triton X-100 (cat #T9284, Sigma-Aldrich) and phalloidin-Alexa 488 (1:100) (A12379, Thermo Fischer Scientific). Images of scratches at *t*_0_ and *t*_24_ were acquired using a high throughput screening microscope (Arrayscan XTI, Thermo Fisher Scientific). For each well, 9 adjacent fields of view were captured and then stitched together as a montage image using a 4x or 20x objective. Images of scratches were analysed using image analysis software (MetaMorph, Molecular Devices, Sa Jose, CA). The analysis protocol consisted of several processing steps - images were smoothened and a filter was applied to create a binary image that separates cells from background, and scratch areas were then measured. Resulting data were used to calculate migration area of cells according to the following formula: Migration area = scratch area in *t*_0_ plate - scratch area in *t*_24_ plate.

### Tube formation assay

LECs were plated in black-walled 96-well imaging plates (Costar ®, Corning) at 1 x 10^4^ cells per well, and transfected with miR mimics or antagomiRs. After 24 h, cells were serum starved, and the rest of the assay was performed 48 h post-transfection. To prepare a 10 ml overlay collagen mix (1 mg/ml collagen), 50 μl of 1 N sodium hydroxide (Sigma-Aldrich), 3.33 ml of 10 x PBS, 950 ml Milli-Q water, 2 ml of bovine collagen I (5 mg/ml) (Gibco), and 6.67 ml serum starvation medium were mixed on ice. The collagen mix (100 μl) was carefully overlaid on top of the cell monolayer to avoid bubbles. Plates were incubated at 37°C in a humidified atmosphere of 5% CO_2_ for 15–30 min to allow collagen to polymerise and form a matrix, after which cells were treated with growth factors for 8 h by pouring serum starvation solution plus growth factors on top of the collagen mix. Cells were then washed once with PBS and fixed with 1% paraformaldehyde (PFA) in PBS for 2 h at room temperature. Cells were then washed in PBS, permeabilised and stained using a PBS solution containing 2% BSA, 0.2% Triton X-100 and phalloidin Alexa 488 (1:100) (A12379, Thermo Fisher Scientific). Images were acquired using a high throughput screening microscope (Arrayscan XTI, Thermo Fisher Scientific) with a 4x or 20x objective. For each well, nine adjacent fields of view were captured and then stitched together as a montage image. Images were processed with MetaMorph image analysis software using an in-built tube formation assay module which measured length and breadth of tubes, number of tubes, number of nodes and number of branch-points.

### Tissue processing and sectioning

All excised tissue specimens were placed in 5 ml of 4% paraformaldehyde solution (cat # C004-1L, 16% Paraformaldehyde aqueous solution, ProSciTech Pty Ltd, Kirwan, Australia), and incubated at 4°C overnight. The next day, tissues were washed with PBS three times and incubated in 12.5% EDTA decalcification solution (Chem-Supply, Gillman, Australia) for 14 days, with the solution being changed every day. Tissues were then removed from EDTA solution, washed with PBS three times and stored in 70% ethanol for at least 24 h. Tissues were then dehydrated in 95% ethanol for 45 min, followed by three incubations in 100% ethanol (45 min each). Tissues were then incubated three times in xylene (45 min each), and embedded in paraffin wax. Formalin-fixed paraffin-embedded tissue blocks were cut into sections 4 μm thick, which were placed onto glass slides coated with 3-aminopropyltriethoxysilane (Sigma-Aldrich) for *in situ* hybridization and immunohistochemistry.

### *In situ* hybridization

Formalin-fixed paraffin-embedded tissues were used for *in situ* hybridization (ISH) employing locked nucleic acid (LNA) probes labelled with digoxigenin (DIG) at both the 5′- and 3’-ends (Qiagen). ISH was performed with probes specific for miR-132-3p (10 nM) or U6 snRNA (2 nM), or with a “scrambled” negative control probe (10 nM), according to manufacturer’s instructions. The scrambled control probe consisted of a random sequence that was not complementary to any known miRNA. Tissue sections were deparaffinized in histolene, rehydrated and treated with proteinase K (15 μg/ml) at 37°C for 10 min. DIG-labelled probes were denatured at 94°C for 5 min. Tissue sections were then coated with probes diluted in hybridization buffer, coverslips applied to avoid evaporation and slides incubated at 56°C for h in a hybridization oven (HybEZ, Oven, Advanced Cell Diagnostics, Newark, CA). Stringent washes were then performed at 56°C: once with 5 x SSC, twice with 1 x SSC, and three times with 0.2 x SSC buffers. Each washing step was carried out for 5 min. Slides were then washed in PBS for 5 min at room temperature. Tissue sections were blocked by incubation with Antibody blocking solution (see Table 2) in a humidified chamber for 15 min. Anti-Digoxigenin-AP Fab fragments (Cat # 11093274910, Roche, Basel, Switzerland) were diluted 1:800 in Antibody Dilutant Solution (Table 2), applied to tissue sections and incubated for 1 h at room temperature in a humidified chamber. Slides were washed three times with PBST for 3 min and tissue sections incubated with AP substrate (Table 2) for 2 h at 30°C. The reaction was stopped by washing slides with KTBT buffer (Table 2) twice for 5 min. Slides were then washed twice in Milli-Q water for 1 min, and tissue sections counterstained with Nuclear Fast Red (N8002, Sigma-Aldrich) for 10 min before carefully rinsing in running tap water for 10 min. Tissue sections were then dehydrated and mounted with Eukitt mounting medium (cat # 03989, Sigma-Aldrich) and coverslips. Images were acquired using an Olympus BX61 microscope and a SPOT Model 25.4 2 Mp Slider digital camera (Diagnostic Instrument Inc. Sterling Heights, MI) using SPOT Software Version 5.0.

**Table 2:**
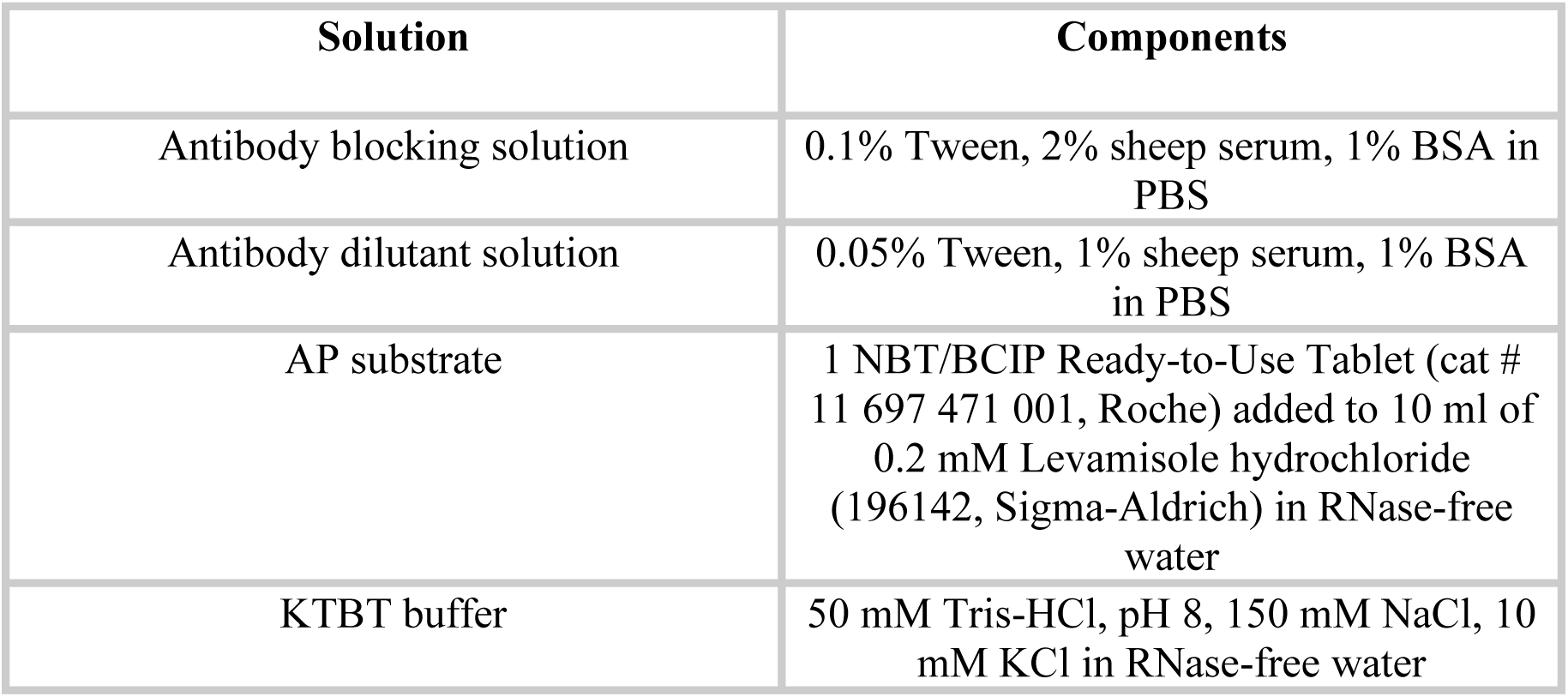
Solutions used for ISH.

### Immunohistochemistry

Immunohistochemical staining was performed for human or mouse podoplanin, human VEGF-D and mouse LYVE1 (see Table 3 for primary antibodies). Formalin-fixed, paraffin-embedded tissue sections were deparaffinized in histolene and rehydrated. For all antigens, slides were incubated in 0.1 M citrate buffer pH 6.0 Target Retrieval Solution (cat # S1699, Dako, Productionsvej, Denmark), and antigen retrieval was performed by heating in a pressure cooker (Dako) at 125°C for 3 min. Slides were left to cool at room temperature for 30 min and washed with PBS three times for 5 min. After antigen retrieval, blocking of endogenous peroxidase activity was performed by incubating tissue sections in 3% H_2_O_2_ in methanol at room temperature for 20 min, followed by rinsing with PBS three times for 5 min. Sections were blocked by incubation with Tris-NaCl blocking buffer (TNB) (0.1 M Tris-HCl, pH 7.5, 150 mM NaCl, and 0.5% w/v blocking reagent, PerkinElmer) for analysis of Podoplanin, or with serum-free protein block (cat # X0909, Dako) for analysis of VEGF-D or LYVE-1 in a humidifying chamber for 1 h. Tissue sections were incubated with the appropriate primary antibody diluted in TNB (for podoplanin) or Antibody Diluent (cat # S0809, Dako) (for VEGF-D or LYVE 1) in a humidified chamber at 4°C overnight. The following day, slides were washed with TNT buffer three times for 5 min and then incubated at room temperature for 1 h with secondary antibodies as follows: (i) For mouse podoplanin, a biotinylated anti-hamster secondary antibody raised in goat (Vector Laboratories, Burlingame, CA) diluted 1:300 in TNB was used; (ii) For human podoplanin and human VEGF-D, a biotinylated anti-mouse secondary antibody raised in goat (Vector Laboratories) diluted 1:200 in Antibody Diluent (Dako) was used; (iii) For mouse LYVE1, a biotinylated anti-rabbit secondary antibody raised in goat (Vector Laboratories) diluted 1:300 in Antibody Diluent (Dako) was used. Slides were then washed with TNT buffer three times for 5 min, before peroxidase staining was conducted for 30 min with a peroxidase staining kit (VECTASTAIN® ABC Reagent, cat # PK-4000, Vector Laboratories) as described by the manufacturer. Slides were then incubated with 3,3’-diaminobenzidine working solution (SK-4100, Vector Laboratories) for 5 to 10 min, as described by the manufacturer, to visualise colour change. The colour reaction was stopped by immersing slides in Milli-Q water for 5 min. Finally, tissue sections were cover slipped with AquaMount (Merck) and left to dry. Images were acquired using an Olympus BX61 microscope and a SPOT Model 25.4 2 Mp Slider digital camera (Diagnostic Instrument Inc.) using SPOT Software Version 5.0. Concentration-matched negative control immunostaining was carried out in all experiments using affinity-purified hamster IgG (400912, Biolegend, San Diego, CA), affinity-purified mouse IgG (sc-2025, Santa Cruz Biotechnology, Dallas, TX) or affinity-purified rabbit IgG (ab37415, Abcam, Cambridge, UK).

**Table 3:** Primary antibodies used for immunohistochemistry.

| Target of antibody | Species of antibody | Type of antibody | Source | Working concentration |
| --- | --- | --- | --- | --- |
| Mouse Podoplanin | Hamster | Monoclonal | Fitzgerald (Cat # 10R-P155A) | 1:1000 |
| Human Podoplanin | Mouse | Monoclonal | GenWay (Cat # GWB-9E8999) | 1:50 |
| Human VEGF-D | Mouse | Monoclonal | VD1 antibody (72) | 35 µg/ml |
| Mouse LYVE1 | Rabbit | Polyclonal | Fitzgerald (Cat # 70R-LR003) | 3 µg/ml |

### Mice

SCID/NOD/gamma (NSG) mice, 6-12 weeks old, were obtained from the Animal Care and Use Program of the Peter MacCallum Cancer Centre (Parkville, VIC, Australia). Mouse experiments were in accordance with the “Australian code for the care and use of animals for scientific purposes”, 8th edition, and were approved by the Peter MacCallum Cancer Centre Animal Experimentation Ethics Committee.

### Ear lymphangiogenesis assay

Mice were anaesthetised by intraperitoneal injection of 20 ml/kg ketamine/xylazine (prepared by combining 0.5 ml of Ketamine (Ketavet 100, Parnell Labs, Alexandria, Australia), 0.5 ml of Xylazine (Ilium Xylazine, Troy Labs, Smithfield, Australia) and 9 ml sterile saline)), and lubricating eye gel (Genteal 0.3%, Alcon Laboratories, Australia) was applied to both eyes to prevent eye dryness. Prior to intradermal ear injections, growth factors, miR mimics and antagomirs were mixed with Matrigel (Matrigel Basement membrane matrix, growth factor-reduced, Phenol Red-Free, ldEV-Free, Cat # FAL356231, BD, Franklin Lakes, NJ). The ear lobes of mice were stretched over the middle finger and a needle (1 ml insulin syringe, 29G) was inserted with an up-facing bevel at an angle of approximately 20 degrees. The dermis was penetrated for approximately 2 mm in length and 20 μl of Matrigel mixture was injected slowly with regular pressure. After injections, mice were housed individually to eliminate the risk of other mice disturbing treated ears. Cages containing injected mice were left on a heat pad for 2 h to allow mice to recover. At the end of experiments (7 days after injection) mice were killed by asphyxiation with CO_2_, and ears were harvested for immunohistochemical analysis.

Ears were fixed in 4% PFA in PBS overnight at 4°C on an orbital shaker. The following day, ears were washed twice in PBS and kept in a solution of 0.3% Triton X-100, 0.05% sodium azide in PBS until dissection. Prior to dissections, ears were carefully shaved with a multi-blade razor, and the frontal side of the ear was then separated from the dorsal side using forceps. Ears were then washed extensively by vortexing in wash solution (0.3% Triton X-100 in PBS). The cartilage was then peeled from the internal part of the frontal side of the ear and another round of washing was performed (by vortexing in wash solution). The fat was then scraped from the internal part of the frontal side of the ear using curved forceps. Ears were then washed for the last time (by vortexing in wash solution) and pinned on a silicone-based matrix (cat # 761028, Sylgard, Sigma-Aldrich) in a 6-well plate. Wash solution was poured into the wells and ears were kept in this solution until whole-mount immunostaining.

Sample sizes for this assay were based on previous in-house experience showing that two independent experiments, each with study group sizes in the range of 8-10, were appropriate from the perspectives of feasibility and statistical comparison. Quantitative analyses for this assay were conducted in a blinded fashion so the investigator was unaware of which sample was from which study group.

### Whole-mount immunohistochemistry on mouse ears

Mouse ears were blocked overnight at 4°C on an orbital shaker in blocking solution (5% goat serum, 0.2% BSA, 0.05% sodium azide and 0.3% Triton X-100 in PBS) and primary antibody treatment was then performed. Ears were stained with LYVE1 antibody (rabbit anti-mouse LYVE1 antibody, cat # 70R-LR003, Fitzgerald, North Acton, MA), diluted 1:1,000 in PBS, to identify lymphatic vessels. This primary antibody was incubated with ears overnight at 4°C on an orbital shaker. Ears were then washed three times (each time for 1 h at room temperature) in washing solution (0.3% Triton X-100 in PBS), and were stored overnight at room temperature in 0.3% Triton X-100, 0.05% sodium azide in PBS. Next day, ears were transferred into a solution containing secondary goat anti-rabbit 488 antibody (cat # A11034, Invitrogen) diluted 1:250 in 0.3% Triton in PBS, and incubated overnight at 4°C on an orbital shaker, protected from light. The next day, ears were washed three times (each time for 1 h at room temperature) in washing solution (0.3% Triton X-100 in PBS) and then stored overnight at room temperature in 0.3% Triton X-100, 0.05% sodium azide in PBS. On the following day, ears were vigorously vortexed for 1 min in a washing solution (0.3% Triton X-100 in PBS) and mounted using aqueous anti-fade mounting medium (Vectashield, cat # H1000, Vector Laboratories) onto slides on which frames (In situ frames, cat # 0030127-536, Eppendorf, Hamburg, Germany) had been applied. Coverslips were then applied and ears imaged during the following week using a confocal microscope (FV3000 Olympus, Tokyo, Japan). For each ear, four fields surrounding the Matrigel plug were imaged and analysed using MetaMorph and AngioTool software (73).

### Argonaute HITS-CLIP

In broad terms, the Ago-HITS-CLIP method was performed as described previously (74) with the following major modifications: after cross-linking, cells were lysed on the plate, and both 3’ and 5’ linkers were ligated to the Ago-bound RNAs prior to elution from the IP beads. More specifically, LECs were transfected in 10 cm cell culture dishes with 20 nM miRVana mimic (miR-132 or negative control) using RNAiMAX (Invitrogen); two technical replicates each of two biological replicates i.e. four samples per condition. After 24 h, cells were rinsed with ice-cold PBS and UV irradiated with 800 mJ/cm^2^, 254 nm, in ice-cold PBS using a UV Stratalinker (Stratagene). Cells were lysed with 1 X PXL (1 X PBS, 0.1% SDS, 0.5% deoxycholate, 0.5% Igepal) containing EDTA-free Complete protease inhibitor cocktail (PIC; Roche). DNA was digested with 5 μl Turbo DNAse (Ambion AM2238) at 37°C for 10 min. RNA was partially digested with RNase 1 (Ambion AM2295) by adding 10 μl of 1:75 diluted RNase 1 per 1 ml of lysate at 37°C for 5 min. Lysates were centrifuged at 55,000 g for 22 min at 4°C and supernatant transferred to a fresh tube. AGO-RNA complexes were immunoprecipitated using mouse IgA2 monoclonal anti-Ago2 antibody 4F9 (75) (hybridoma sourced from University of Florida ICBR). Antibodies (15 μg) were conjugated to 20 μl protein L Dynabeads (ThermoFisher, 88849) before resuspending the washed beads with 1 ml of prepared lysate at ∼800 μg/ml and rotating for 2 h at 4°C. Bound AGO-RNA complexes were washed twice each consecutively with ice cold 1 X PXL, 5 X PXL (5 X PBS, 0.1% SDS, 0.5% sodium deoxycholate, 0.5% Igepal), and 1 X PNK (50 mM Tris-Cl pH 7.5, 10 mM MgCl2, and 0.5% Igepal). Beads were first treated with T4 PNK (NEB, M0201L; 20 U in 80 μl reaction volume) in the absence of ATP (37°C, 850 rpm for 20 min) to dephosphorylate 3’ RNA ends followed by washes with 1 X PNK, 5 X PXL, and two washes with 1 X PNK at 4°C. The 3’ preadenylated linker (NEBNext 3’SR adaptor for Illumina; /5rApp/AGA TCG GAA GAG CAC ACG TCT /3AmMO/) was ligated to the RNA fragments on bead using T4 RNA ligase 2 truncated KQ (NEB M0373) at 16°C, overnight with shaking. Beads were washed consecutively with ice cold 1 X PXL, 5 X PXL, and twice with 1 X PNK. Bound RNAs were then labelled with P32 γ-ATP using T4 PNK, 25 min at 37°C, followed by addition of 2.5 µM ATP, 5 min at 37°C. Beads were washed twice each with ice-cold 1 X PNK+EGTA (50 mM Tris-HCL pH7.5, 20 mM EGTA, 0.5% Igepal) and 1 X PNK. The 5’ RNA linker (5’-blocked and containing a 10 nt UMI (/5AmMC6/GUUCAGAGUUCUACAGUCCGACGAUCNNNNNNNNNN3’) was ligated to the RNA fragments on bead using T4 RNA ligase (NEB M0437) for 2 h at 24°C with shaking. Beads were washed 3 times with ice-cold 1 X PNK+EGTA.

AGO-RNA complexes were eluted with 40 μl 1 X Bolt LDS sample buffer (ThermoFisher), 1% β-mercaptoethanol at 70°C for 10 min. Samples were separated through Bolt 8% Bis-tris Plus gels (ThermoFisher) using BOLT MOPS SDS running buffer at 165 V for 75 min. Complexes were then transferred to nitrocellulose (Schleicher&Schuell, BA-85) by wet transfer using 1 X BOLT transfer buffer with 10% methanol. Filters were placed on a phosphor screen and exposed using a Typhoon imager (GE). 130-180 kDa regions (corresponding to RNA tags > 25 nt) were excised from the nitrocellulose. RNA was extracted by proteinase K digestion (2 mg/mL proteinase K, 100 mM Tris-HCl pH 7.5, 50 mM NaCl, 10 mM EDTA, 0.2% SDS) at 50°C for 60 min on a Thermomixer (1,200 rpm) followed by extraction with acid phenol (ThermoFisher, AM9712) and precipitation with 1:1 isopropanol:ethanol. RNA was pelleted by centrifugation then separated on a 15% denaturing polyacrylamide gel (1:19 acrylamide, 1 X TBE, 7 M urea). The wet gel was wrapped in plastic wrap and exposed to a phosphor screen and imaged using a Typhoon. Gel slices were cut corresponding to the expected size of the cross-linked RNA (70 – 280 nt) and eluted by the “crush and soak” method as previously described (76). Reverse transcription was performed using SR-RT primer (IDT, AGACGTGTGCTCTTCCGATCT) with SuperScript IV. Products were amplified for 12 cycles using a common forward primer (NEBNext SR primer for Illumina) and barcoded reverse primers for each sample (NEBNext Index primers for Illumina). PCR products were purified using 1.8 volumes of Axygen AxyPrep magnetic beads (MAG-PCR-CL), separated on an 8% acrylamide (19:1), 7 M urea, TBE semi-denaturing gel, stained with SYBR Gold nucleic acid gel stain (ThermoFisher) and imaged on a ChemiDoc (BioRad). Products corresponding to an insert size of ∼25 – 70 nt were excised from the gel and extracted by the “crush and soak” method as above. Library quality and quantity was assessed by Bioanalyzer (Agilent) and qPCR, pooled and sequenced on an Illumina NextSeq 500 (1 x 85bp).

### Bioinformatic analyses

#### Small RNA sequencing

To perform small RNA sequencing, RNA quality was analysed with Agilent 2100 Bioanalyzer and Agilent RNA 6000 Nano kit to confirm RIN values were above 7, and RNA concentration was determined with Qubit High Sensitivity RNA kit (Thermo Fisher). Total RNA (1 μg) was then used for small RNA library preparation with the NEBNext Multiplex Small RNA Library Prep Set for Illumina (Set 2) according to manufacturer’s guidelines. Barcoded libraries were amplified with 15 cycles of PCR and size selected via Pippin prep with 3% DF Marker F gel. Library sizes around 150 bp were confirmed using an Agilent 2100 Bioanalyzer and the Agilent High Sensitivity DNA kit. Yields of libraries were confirmed with the Qubit dsDNA High Sensitivity assay. Libraries were then diluted to generate 1 nM stocks, whereupon multiple libraries were pooled together in equimolar ratios. Libraries were then sequenced on a single Illumina HiSeq 2000 run, generating single-end, 50 bp reads with an average of ∼19 million reads per library. FASTQ files were analysed at various stages for quality and content using FastQC v0.11.2 (77). Raw reads were adapter trimmed and filtered using cutadapt v1.3 (78) using an adapter sequence of AGATCGGAAGAGCACACGTCTGAACTCCAGTCAC, minimum length of 18, error rate of 0.2, and overlap of 5. The resulting reads were mapped against the human reference genome (build hg19) using the BWA v0.7.9a-r786 alignment algorithm (79) with default parameters. Annotation obtained from miRBase (www.mirbase.org; (80)) was used to obtain miR-level read counts using htseq-count (81) and read counts were normalised to reads per million (RPM) using the total number of reads mapping to microRNAs.

#### HITS-CLIP

HITS-CLIP libraries were prepared for LEC cells transfected with miR-132 mimic or a negative control mimic, consisting of two biological replicates which were each separated into two technical replicates and sequenced to produce 85 bp single-ended reads with average sequencing depth of ∼22 million reads. FASTQ files were analysed at various stages for quality and content with FastQC v0.11.5 (77) and raw reads were adapter trimmed and filtered using cutadapt v1.6 (78) with an adapter sequence of AGATCGGAAGAGCACACGTCTGAACTCCAGTCA, error rate of 0.2, overlap of 5 and minimum length of 18. Reads derived from PCR duplication were collapsed using Unique Molecular Identifiers (UMIs) with UMI-tools (v0.5.3; (82)) by first using the ‘extract’ method with default parameters to cut the 10 bp UMIs from the 3’ end of the reads allowing an edit distance of 1. To address cases where 5’ adapters had concatemerized during preparation of the libraries, a second round of adapter trimming was performed to remove the 5’ adapter using cutadapt with the same parameters as above but an adapter sequence of GTTCAGAGTTCTACAGTCCGA. Filtered reads were mapped against the human reference genome (hg19) using the Tophat2 alignment algorithm (version 2.1.1 with default parameters) (83), returning an average alignment rate of ∼45%. Subsequently, UMIs were used to collapse PCR duplicate reads using the UMI-tools ‘dedup’ method with default parameters. To identify enriched regions of the genome, technical and biological replicates were pooled using the Picard Tools function MergeSamFiles (84) and quality filtered using samtools (-q 5) (85). Peak calling was then performed separately for each strand using MACS2 peak caller (version 2.1.1) (86) with the negative control mimic sample as control (settings: -f BAM -g hs --keep-dup all --nomodel --shift -15 --extsize 50 -B --call-summits --slocal 0 --llocal 0 --fe-cutoff 2 -q 0.05) and the resulting peak files from each strand were merged. HITS-CLIP peaks and alignments were visualized and interrogated using the Integrative Genomics Viewer v2.8.0 (87) and the Targetscan (v7.2) “Total context++ score” was used to identify predicted targets of miR-132. Homer (88) was used to identify peaks located in 3’ UTRs and annotate peaks with positions of DNA sequences matching a 6 nt portion of the miR-132 seed region (AACAGT). Predicted mRNA targets of miR-132 from Ago HITS-CLIP were subjected to gene ontology analysis with QuickGo (https://www.ebi.ac.uk/QuickGO/) and pathway analysis with The Database for Annotation, Visualization and Integrated Discovery (DAVID) using default parameters. Sequencing read and peak files for the HITS-CLIP dataset are available from the NCBI Gene Expression Omnibus under the accession GSE190833 (for Reviewer access go to https://www.ncbi.nlm.nih.gov/geo/query/acc.cgi?acc=GSE190833 and enter token kfwtowawhjizhub into the box).

### Human tissues

Use of human tissue was approved by the Human Ethics Committee of the Peter MacCallum Cancer Centre (approval number 10/16).

### Statistical analyses

Each experiment involved two, or most frequently three, independent biological replicates with any exceptions stated. All error bars in graphs represent standard error of the mean (SEM) unless stated otherwise. All statistical analyses were performed using a statistics software package (Prism 8 for macOS; GraphPad Software). Statistical significance was evaluated by Student’s t-test or one-way analysis of variance (ANOVA) with Tukey’s method used to correct for multiple comparisons.

## SUPPLEMENTARY MATERIAL

**Supplementary Datafile 1.** List of mRNA targets of miR-132 as identified by Ago HITS-CLIP. Approximately 900 miR-132 target sites are listed involving approximately 500 mRNAs.

## ACKNOWLEDEGMENTS

We thank the following: David Byrne, Elena Takano, Kaushalya Amarasinghe and Jason Li (Peter MacCallum Cancer Centre) for technical assistance, Rae Farnsworth (Peter MacCallum Cancer Centre) for helpful discussion and Biorender (https://biorender.com/) for design of diagrams. This study was supported by Program, Project, Ideas and Investigator Grants, and Senior Research Fellowships, from the National Health and Medical Research Council of Australia, and by funds from the O’Brien Foundation, Stafford Fox Foundation, and Wicking Foundation, Australia, and the Operational Infrastructure Support Program provided by the Victorian Government, Australia. V.A. was supported by a University of Melbourne Postgraduate Scholarship from the University of Melbourne, Australia.

